# The Neuro Bureau ADHD-200 Preprocessed Repository

**DOI:** 10.1101/037044

**Authors:** Pierre Bellec, Carlton Chu, François Chouinard-Decorte, Yassine Benhajali, Daniel S. Margulies, R. Cameron Craddock

**Author notes:** Email addresses:* (Pierre Bellec), (Carlton Chu), (François Chouinard-Decorte), (Yassine Benhajali), (Daniel S. Margulies), (R. Cameron Craddock). URL:* bellec.simexp-lab.org (Pierre Bellec), computational-neuroimaging-lab.org, (R. Cameron Craddock).

## Abstract

In 2011, the “ADHD-200 Global Competition” was held with the aim of identifying biomarkers of attention-deficit/hyperactivity disorder from resting-state functional magnetic resonance imaging (rs-fMRI) and structural MRI (s-MRI) data collected on 973 individuals. Statisticians and computer scientists were potentially the most qualified for the machine learning aspect of the competition, but generally lacked the specialized skills to implement the necessary steps of data preparation for rs-fMRI. Realizing this barrier to entry, the Neuro Bureau prospectively collaborated with all competitors by preprocessing the data and sharing these results at the Neuroimaging Informatics Tools and Resources Clearinghouse (NITRC) (http://www.nitrc.org/frs/?group_id=383). This “ADHD-200 Preprocessed” release included multiple analytical pipelines to cater to different philosophies of data analysis. The processed derivatives included denoised and registered 4D fMRI volumes, regional time series extracted from brain parcellations, maps of 10 intrinsic connectivity networks, fractional amplitude of low frequency fluctuation, and regional homogeneity, along with grey matter density maps. The data was used by several teams who competed in the ADHD-200 Global Competition, including the winning entry by a group of biostaticians. To the best of our knowledge, the ADHD-200 Preprocessed release was the first large public resource of preprocessed resting-state fMRI and structural MRI data, and remains to this day the only resource featuring a battery of alternative processing paths.

## 1. Introduction

In 2011, the “ADHD-200 Global Competition” was held with the aim of engaging researchers from a variety of analytical backgrounds to identify biomarkers of attention-deficit/hyperactivity disorder (ADHD) from resting-state functional magnetic resonance imaging (rs-fMRI) and structural MRI (s-MRI) data [1]. The competition made use of the “ADHD-200 Sample” data collection that was aggregated from eight independent sites and shared through the Intenational Neuroimaging Datasharing Initiative (INDI) [2]. The data includes rs-fMRI, structural MRI (s-MRI), and basic phenotypic information for 973 individuals: some typically-developing controls (TDC) and patients diagnosed with ADHD [1]. Competitors were given five and a half months to optimize a classification algorithm on training data (776 individuals) and submit their predicted clinical labels on test data for which diagnostic information was withheld. The competition data was distributed in a raw form and, before any analysis could begin, the images had to be preprocessed to make them comparable across individuals and reduce noise. These preprocessing steps present a significant hurdle for would-be competitors who do not have the specialist knowledge of neuroimaging methods, or access to high performance computing resources. Realizing this barrier to entry, the Neuro Bureau, a non-profit organization aimed at facilitating open science grassroots initiatives^1^, prospectively collaborated with all competitors by preprocessing the data and sharing these results.

The “ADHD-200 Preprocessed” is a repository of preprocessed rs-fMRI and s-MRI data along with statistical derivatives from the ADHD-200 Sample. Rather than favoring a specific processing strategy, we followed a pluralistic approach by preprocessing the data using multiple pipelines (called “Athena”, “Burner”, and “NIAK”) that differed in the toolsets used, the philosophy motivating choices of algorithms and parameters, and the statistical derivatives calculated. The Athena pipeline processed rs-fMRI and s-MRI images using a combination of AFNI [3] and FSL [4] neuroimaging toolkits. The Burner pipeline used SPM8 [5] to process s-MRI data for voxel-based morphometry. The NIAK pipeline processed rs-fMRI and s-MRI using the NeuroImaging Analysis Kit [6].

## 2. Organization and access to the repository

The ADHD-200 Preprocessed data was released in 2011 and can be downloaded from NITRC^2^. No data usage agreement is required to access or download the data, the only requirement is registering for a free NITRC account. This registration enables downloads to be tracked for usage statistics and users to be contacted in the event that errors are found in the dataset. The ADHD-200 Sample allows unrestricted data usage for non-commercial research purposes provided that the specific datasets included in an analysis be cited appropriately and that their funding sources be acknowledged^3^. There are no more restrictions placed on the preprocessed data or derivatives other than the request that the ADHD-200 Preprocessed Initiative is cited appropriately and that the specific pipeline is acknowledged in publications using the data. A forum is available on the Neuro Bureau’s NITRC project page for users to ask questions or report problems^4^. Questions regarding data acquisition or phenotypic variables should be directed to INDI’s support forum^5^

## 3. Contents of the repository

The ADHD-200 Preprocessed repository contains preprocessed outputs and derivatives for data from the ADHD-200 Sample, which includes 973 individuals (352 F) between the ages of 7 and 27 aggregated from 17 different studies conducted across 8 different sites (for a breakdown of age and sex by diagnosis, see Table 1). For each individual, phenotypic data includes sex, age, handedness, ADHD diagnosis (585 TDC, 362 ADHD, 26 with diagnosis unavailable), ADHD subtype (ADHD-combined, ADHD-inattentive, ADHD-hyperactive/impulsive), one of three different measures of ADHD severity, one of five measures of intelligence, co-morbid diagnoses, and whether or not they have used medication to treat their symptoms [1]. Imaging data for each individual includes one or more T1-weighted high-resolution s-MRI scan (s-MRI) and one or more rs-fMRI scan. The majority of data was acquired during a single imaging session, although a second session is available for 15 individuals from the Washington University at Saint Louis (WUSTL) site. There is a substantial amount of variation in data acquisition procedures across sites including the type of MRI system and scanning parameters, the length of the rs-fMRI scans, and the instructions given to participants prior to the scan (see Table 2 and Table 3).

**Table 1:**
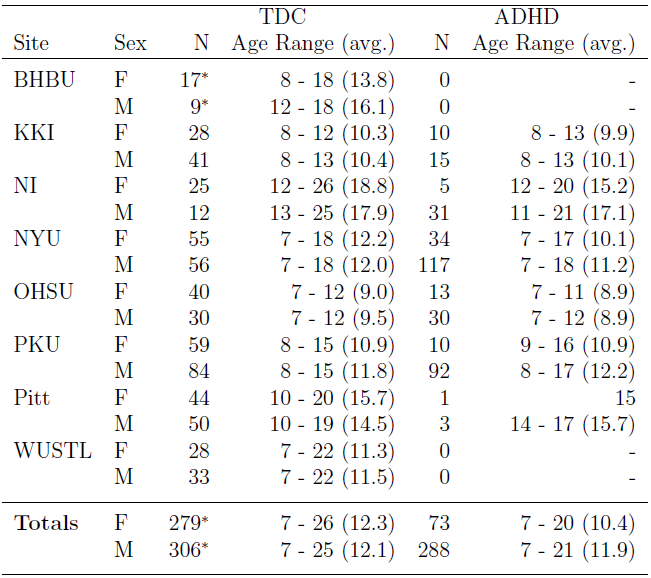
**ADHD-200 participants by site**. BHBU: Bradley Hospital/ Brown University, KKI: Kennedy Krieger Institute, NI: NeuroIMAGE sample, NYU: New York University Child Study Center, OHSU: Oregon Health Sciences University, PKU: Peking University, Pitt: University of Pittsburgh, WUSTL: Washington University at Saint Louis, avg.: average. *Diagnostic labels are currently not available for BHBU, they have been listed as TDC in the table, but not included in the totals.

**Table 2:**
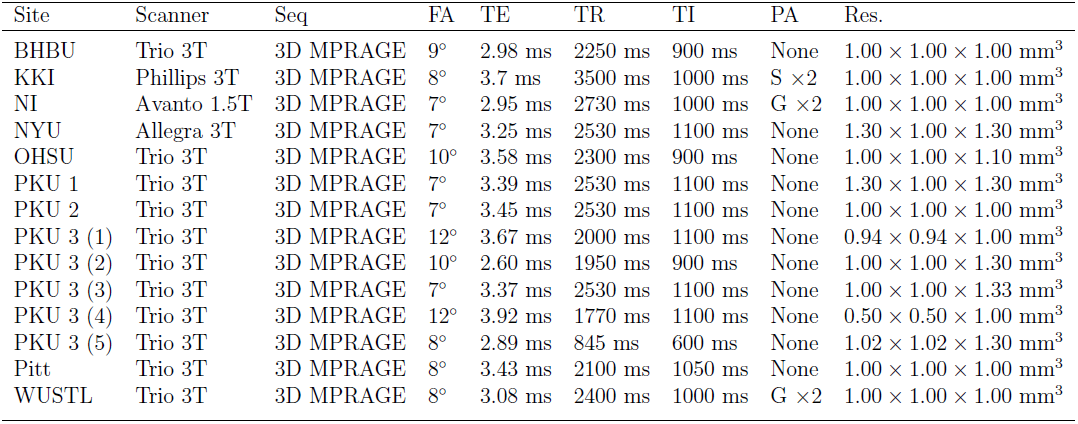
**Structural MRI acquisition parameters by site**. Seq: imaging sequence, FA: flip angle, TE: echo time, TR: repetition time, TI: inversion recovery delay, PA: parallel acquisition, Res: voxel resolution, BHBU: Bradley Hospital/ Brown University, KKI: Kennedy Krieger Institute, NI: NeuroIMAGE sample, NYU: New York University Child Study Center, OHSU: Oregon Health Sciences University, PKU: Peking University, Pitt: University of Pittsburgh, WUSTL: Washington University at Saint Louis, Trio: Siemens TIM Trio 3T, Allegra: Siemens Allegra, Avanto: Siemens Avanto, MPRAGE: magnetization prepared rapid gradient echo, S: sensitivity encoding (SENSE), G: generalized auto-calibrating partially parallel acquisition (GRAPPA)

**Table 3:** **Resting state fMRI acquisition parameters by site**. Seq: imaging sequence, FA: flip angle, TE: echo time, TR: repetition time, PA: parallel acquisition, N_*slc*_: number of slices, Th.: slice thickness, Slc. Acq.: slice acquisition order, N_*TR*_: number of measurements (TRs), BHBU: Bradley Hospital/ Brown University, KKI: Kennedy Krieger Institute, NI: NeuroIMAGE sample, NYU: New York University Child Study Center, OHSU: Oregon Health Sciences University, PKU: Peking University, Pitt: University of Pittsburgh, Pitt 2: U. Pitt. parameters used for acquiring the testing data, WUSTL: Washington University at Saint Louis, EPI: echo planar imaging, PACE: Prospective Acquisition CorrEction (EPI with prospective motion correction), S: sensitivity encoding (SENSE), G: generalized autocalibrating partially parallel acquisition (GRAPPA), int+: slices were acquired interleaved ascending, seq+: slices were acquired sequentially ascending, var.: the number of measurements varies across datasets, fixate: participants were asked to keep their eyes open and fixate on an image, closed: participants were asked to keep their eyes closed, open: participants were asked to keep their eyes open.

| Site | Seq | FA | TE | TR | PA | $N_{slc}$ | Th. | Slc. Acq. | Resolution | $N_{TR}$ | Instructions |
| --- | --- | --- | --- | --- | --- | --- | --- | --- | --- | --- | --- |
| BHBU | PACE | 90° | 25 ms | 2000 ms | None | 35 | 3 mm | int+ | $3.0 \times 3.0 \text{ mm}^2$ | 256 | fixate |
| KKI | EPI | 75° | 30 ms | 2500 ms | $S \times 3$ | 47 | 3 mm | seq+ | $3.0 \times 3.0 \text{ mm}^2$ | 128 | fixate |
| NI | EPI | 80° | 40 ms | 1960 ms | $G \times 2$ | 37 | 3 mm | int+ | $3.5 \times 3.5 \text{ mm}^2$ | 266 | eyes closed |
| NYU | EPI | 90° | 15 ms | 2000 ms | None | 33 | 4 mm | int+ | $3.0 \times 3.0 \text{ mm}^2$ | 180 | eyes closed |
| OHSU | EPI | 90° | 30 ms | 2500 ms | None | 36 | 3.8 mm | int+ | $3.8 \times 3.8 \text{ mm}^2$ | 82 | fixate |
| PKU 1 | EPI | 90° | 30 ms | 2000 ms | None | 33 | 4.2 mm | int+ | $3.1 \times 3.1 \text{ mm}^2$ | 240 | closed or fixate |
| PKU 2 | EPI | 90° | 30 ms | 2000 ms | None | 33 | 3.6 mm | int+ | $3.1 \times 3.1 \text{ mm}^2$ | 240 | closed or fixate |
| PKU 3 | EPI | 90° | 30 ms | 2000 ms | None | 30 | 4.5 mm | int+ | $3.44 \times 3.44 \text{ mm}^2$ | 240 | closed or fixate |
| Pitt | EPI | 70° | 29 ms | 1500 ms | $G \times 2$ | 29 | 4.0 mm | int+ | $3.1 \times 3.1 \text{ mm}^2$ | 200 | open or closed |
| Pitt 2 | EPI | 90° | 30 ms | 3000 ms | None | 46 | 3.5 mm | int+ | $3.8 \times 3.8 \text{ mm}^2$ | 128 | open or closed |
| WUSTL | EPI | 90° | 27 ms | 2500 ms | None | 32 | 4.0 mm | int+ | $4.0 \times 4.0 \text{ mm}^2$ | var. | fixate |

**Table 4:**
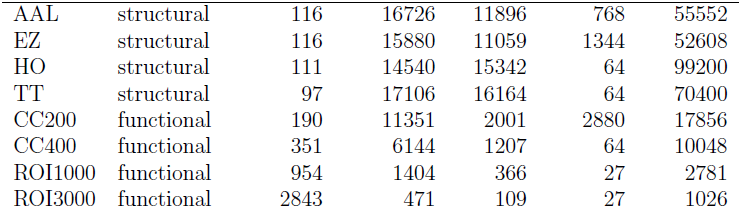
**Summary of the characteristics of all brain parcellations used in the ADHD-200 Preprocessed release**. Sizes for the parcels are reported in mm^3^.

| Name | Type | # parcels | mean size | std size | min size | max size |
| --- | --- | --- | --- | --- | --- | --- |
| AAL | structural | 116 | 16726 | 11896 | 768 | 55552 |
| EZ | structural | 116 | 15880 | 11059 | 1344 | 52608 |
| HO | structural | 111 | 14540 | 15342 | 64 | 99200 |
| TT | structural | 97 | 17106 | 16164 | 64 | 70400 |
| CC200 | functional | 190 | 11351 | 2001 | 2880 | 17856 |
| CC400 | functional | 351 | 6144 | 1207 | 64 | 10048 |
| ROI1000 | functional | 954 | 1404 | 366 | 27 | 2781 |
| ROI3000 | functional | 2843 | 471 | 109 | 27 | 1026 |

Nearly all of the imaging data from the ADHD-200 Sample was included in the preprocessing effort, though some individuals were excluded for poor quality or missing data^6^. The results of the preprocessing are made available as a collection of compressed tar files that are organized by pipeline, sites of data collection, training and test samples, as well as by derivatives. A group-level file containing the phenotypic data is available in comma-separated-values format (.csv).

Shared preprocessed data and extracted features include:

- 3D grey matter density maps suitable for voxel-based morphometry – Athena and Burner (see Figure 1),
- 4D preprocessed resting-state fMRI data including limited intermediaries and quality assessment – Athena and NIAK,
- Average time series for brain regions from structurally defined parcellations – Athena (see Figure 2 and Figure 3),
- Average time series for brain regions for regions defined by functional parcellation – Athena and NIAK (see Figure 2 and Figure 3),
- Spatial maps for 10 intrinsic connectivity networks (ICNs), fractional amplitude of low frequency fluctuations (fALFF), and regional homogeneity (ReHo)-Athena (see Figure 4).

**Figure 1:**
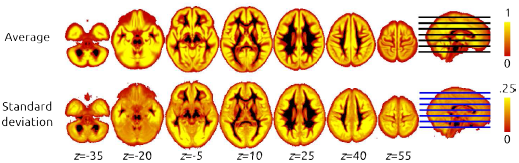
Average and standard deviation of the grey matter density maps generated by the Burner pipeline, for all subjects in the test subsample of ADHD-200 Preprocessed.

**Figure 2:**
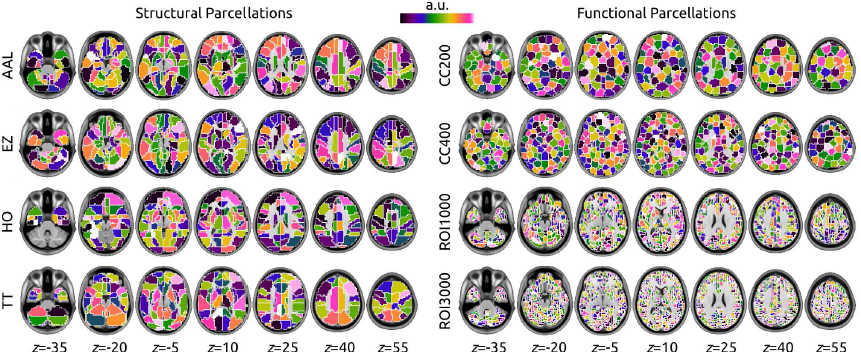
The brain parcellations used to generate regional time series in the NIAK (ROI1000 and ROI3000) and Athena (all other parcellations) pipelines. Each region was randomly assigned to one color in the colormap, and the in-plane outline of regions was painted white at 1 mm resolution.

**Figure 3:**
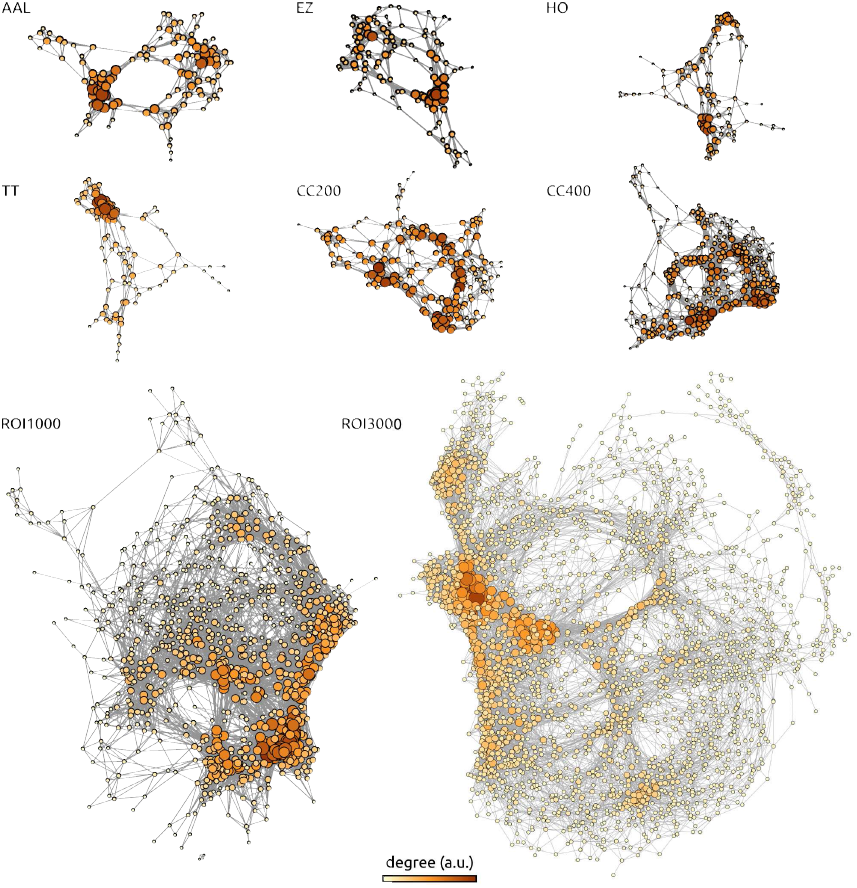
The average functional connectivity matrix (Pearson’s correlation coefficient between regional time series) was generated across all individuals of the KKI site, for all parcellations of the release (see text for details). This matrix was further binarized by retaining connections with an average correlation larger than 0.3. The resulting binary adjacency matrices have been represented with an automated layout generated by Yfan Hu’s multilevel algorithm, as implemented in the Gephi software [7]. The size and color of each node was set proportional to its degree, relative to the min and max inside the graph.

### 3.1 Athena Pipeline

The Athena pipeline^7^ processed rs-fMRI and s-MRI images using a custom BASH script that combined AFNI [3] and FSL [4] neuroimaging toolkits and was run on the Athena computer cluster at Virginia Tech’s Advanced Research Computing center^8^. The processing scripts for each site are distributed in the repository, as well as on Github^9^, along with output log files for each processed dataset.

**Figure 4:**
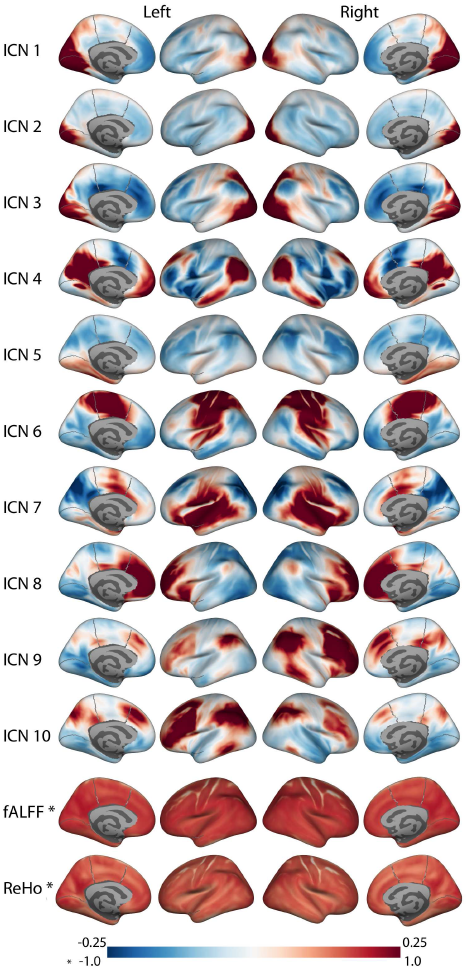
Derivatives from the Athena pipeline including ten intrinsic connectivity networks (ICNs), fractional amplitude of low-frequency fluctuations (fALFF), and regional homogeneity (ReHo). Asterisks (*) in the latter two derivatives denote the difference in colorbar max/min values, as indicated.

#### 3.1.1 Structural processing

Athena’s s-MRI pipeline began with skull-stripping to remove non-brain tissue and background from the images [8] and segmenting the results into white matter (WM), cerebrospinal fluid (CSF), and grey matter (GM) probability maps [9]. A non-linear warp was calculated between the skull-off image and MNI space as represented by the NIHPD 4.5-18.5y age-specific assymmetric template [10] using a two step procedure that calculates a linear transform [11], that is subsequently refined by a non-linear registration procedure [12]. *Shared s-MRI outputs include*: skull-stripped whole-brain images and smoothed (by a 6 mm FWHM Gaussian) and unsmoothed GM density maps in MNI space at 1×1×1 mm^3^ resolution, along with the FSL fNIRT non-linear warp, as compressed NIfTI files (.nii.gz).

#### 3.1.2 Functional processing

*Preprocessing*. Athena’s rs-fMRI pipeline involved removing the first four volumes to allow for magnetization to reach equilibrium, site-specific slice timing correction to the middle slice, re-aligning each volume to the first volume to correct for motion [13], and calculating a linear transform between the mean functional volume and the corresponding s-MRI [11]. The rs-fMRI to s-MRI transform was then combined with the s-MRI to MNI non-linear warp to write the functional data into MNI152 space at 4 × 4 × 4 mm^3^ resolution. Mean WM and CSF signals extracted using the masks calculated during s-MRI processing were included along with 6 head motion parameters and a third-order polynomial in voxelwise nuisance regression models to remove variation due to physiological noise, head motion, and scanner drifts from the time series[14,15]. The resulting denoised time series were band-pass filtered (0.009 Hz < f < 0.08 Hz) to limit the data to the frequencies implicated in resting state functional connectivity [16, 17] and then spatially smoothed with a 6 mm FWHM Gaussian filter. *Shared rs-fMRI outputs include*: denoised rs-fMRI volumes, with and without temporal bandpass filtering, in MNI space (compressed 4D NIfTIs, nii.gz), the mean rs-fMRI image and brain mask in template space (.nii.gz), and six parameter head motion traces (tab-separated values, AFNI .1D files).

*Time series for structurally defined brain areas*. Regional time series were extracted for the automated anatomical labeling (AAL) [18], Eickhoff-Zilles (EZ) [19], Harvard-Oxford (HO) [20–23], and Talairach and Tournoux (TT) [24] parcellations. The EZ parcellation was derived from the max-propagation parcellation distributed with the SPM Anatomy Toolbox^10^ and was transformed into template space using the Colin 27 template (also distributed with the toolbox) as an intermediary. The HO parcellation was constructed from 25% thresholded cortical and subcortical max-propagation parcellations distributed with FSL. The parcellations were bisected into left and right hemispheres at the midline (*x* = 0), ROIs representing left/right WM, left/right GM, left/right CSF and brainstem were removed from the sub-cortical parcellation and then the subcortical and cortical ROIs were combined into a single parcellation. The AAL parcellation distributed with the SPM8 version of the AAL Toolbox^11^ and the TT parcellation distributed with AFNI were coregistered and warped into template space. Each of the structural parcellations were resampled into the functional space using nearest-neighbor interpolation. The average time series within each parcel were extracted from both the filtered and unfiltered data, and are distributed in tab-separated values format (AFNI .1D). Each of the conformed ROI parcellations are available as compressed 3D NIfTI files (.nii.gz).

*Time series for functionally defined parcellations*. The CC200 and CC400 functional brain parcellations were constructed using a two-stage spatially-constrained spectral clustering procedure [25] applied to unfiltered preprocessed rs-fMRI data from a subset (*N* = 650) of the participants in the training dataset. Participants were chosen for inclusion based on registration quality and after excluding participants with more than 3 mm translation or 3 degrees rotations in their motion parameters. To reduce computation time, the clustering was restricted to grey matter using a group GM mask that was constructed by averaging individual GM masks derived from FreeSurfer automated segmentation [26]. Although 200 and 400 ROIs were specified in the functional parcellation procedure, the normalized cut algorithm resulted in 190 and 351 clusters respectfully. Time series were extracted for each parcellation from both the filtered and unfiltered data by averaging the voxel time series contained within each labeled region and are distributed in tab-separated values format (AFNI .1D). CC200 and CC400 brain parcellations are available as compressed 3D NIfTI files (.nii.gz).

*ICN time series and spatial maps*. Time series and spatial maps were derived for 10 group ICNs generated by [27], which were found to be consistent across resting-state datasets and a variety of neuroimaging tasks. Based on these template ICNs, we applied a modified dualregression approach [28] to the unfiltered preprocessed data. A spatial multiple regression was first used to extract time series corresponding to each network. In a second step, each time course was independently correlated with whole-brain time series to generate subject-specific functional connectivity maps for each network. Alternatively, all time series were entered simultaneously into a multiple (temporal) regression, and the regression coefficients associated with each time series constituted the functional connectivity maps. The resulting ICN time series are distributed as tab-separated values (AFNI .1D) files and the spatial maps for both temporal regression approaches are distributed as compressed 4D NIfTI files (.nii>.gz).

*fALFF and ReHo*. Whole brain fALFF maps were generated by dividing the variance of each voxel’s bandpass-filtered time series by the variance of its unfiltered time series [29].ReHo was estimated from the unfiltered data at each voxel by the Kendall’s Coefficient of Concordance [30] between the voxel and its 26 face-, edge-, and corner-touching neighbors. The resulting fALFF and ReHo whole brain maps are distributed as compressed 3D NIfTI files (.nii.gz).

#### 3.1.3 Quality Control

Images were visually inspected and attempts were made to fix gross misregistration errors by hand adjusting the offending images to center them on the anterior commisure and rotate them into rough correspondence with the MNI template. With the exception of a few datasets that were missing data, and one dataset that was corrupted, all of the ADHD-200 Sample was processed and released by Athena, regardless of data quality. This was done to accommodate differing opinions as to what qualifies as usable data, and to provide poor quality data that may be used by others to develop methods that are robust to noise. Files containing the six motion parameters for each rs-fMRI data and anatomical and mean EPI images in template space have been included in the release to enable users to determine high-motion data or poor registrations.

Additional quality metrics derived from the preprocessed s-MRI, GM masks, mean rs-fMRI images, fALFF maps, and FC maps were also included to help with the QC process. For each data type a mean and standard deviation image was calculated from all of the scans (in stereotaxic space) from all of the subjects. These images were used to perform a voxelwise z-score transformation on each data type for each subject (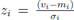). For each image (or map), the number of these z-scores whose absolute value exceeded 3 where summed to generate a quality score. The higher the resulting sum, the larger the number of voxels a image has with |*z*| ≥ 3 and the more likely the images are outliers. Using this metric it is possible to rank images for a more directed search for poor quality scans. These metrics are distributed in participant-specific text files along with the BASH scripts that implement the procedure.

A rigorous visual inspection of registration has also been performed using the same procedures and raters (PB and YB) as the NIAK pipeline. The procedure entailed generating an online report^12^ of registration quality for all subjects. The report includes a visualization of the individual structural images in standard space overlaid onto the corresponding brain template (MNI pediatric), as well as a visualization of the individual mean EPI image overlaid onto the corresponding structural image, both in standard space. These images were visually reviewed, and images with severe misregistraion or quality problems were marked as “fail” (7% failure rate). The outcome of the visual quality assessment along with all available phenotypic variables have been consolidated into a tabular-separated (.tsv) spreadsheet that can be downloaded from NITRC^13^.

### 3.2 Burner Pipeline

The Burner pipeline^14^ used SPM8 [5] to process s-MRI data for voxel-based morphometry [31] style analyses.

#### 3.2.1 Structural processing

Processing began by segmenting s-MRI images into GM and WM probability maps using SPM8’s unified segmentation procedure, which iteratively registers the data to a template and performs tissue classification until both are optimized [32]. Next, SPM8’s DARTEL toolbox [33] was used to register the s-MRI of all participants into a common space using an iterative method. Initially, all WM and GM maps were rigidly aligned, and the initial GM and WM templates were created by averaging all aligned maps. Then, all WM and GM maps were non-linearly registered to the templates. New templates were created after each such iteration of registration. The procedure was repeated six times (i.e. template creation and registration) to generate sharper templates and warping all participant WM and GM maps to the template space. The final (6th iteration) non-linear deformations were applied to each participant’s GM probability maps to transform them into the space of the population average at 1.5 × 1.5 × 1.5 mm^3^ resolution and modulated to conserve the global tissue volumes after normalization. The resulting grey matter density maps are distributed as 3D NIfTI files (.nii).

#### 3.2.2 Quality Control

Stringent quality control was not applied to the data in order to accommodate different opinions on what constitutes poor quality data. Images for four participants were excluded after visual inspection by Dr. Chu because they were determined to be of insufficient quality for further processing.

### 3.3 NIAK Pipeline

The NIAK^15^ is a collection of workflows, implemented in the Pipeline System for Octave and Matlab (PSOM) [34], that perform s-MRI and rs-fMRI processing using a combination of generic medical image processing modules, the MINC tools^16^, and custom Matlab/Octave scripts. The ADHD-200 Sample was processed using NIAK version 0.6.4.1, running on a server of the Canadian Brain Imaging Research Platform (CBRAIN) [35]. The NIAK is distributed as an open-source software under MIT license and the code is available on NITRC^17^ and Github^18^. The processing scripts for ADHD200 are available on github^19^. The log files for execution were included with the derivatives and can be accessed through the PSOM interface^20^.

#### 3.3.1 Structural processing

The NIAK implements a variant of the CIVET pipeline [36]. Each individal s-MRI scan was first corrected for intensity non-uniformities [37] and the brain was extracted using a region growing algorithm [38]. Individual scans were then linearly registered (9 parameters) with the T1 MNI symmetric template [10], restricted to the brain with the previous mask. Note that, by selecting a symmetric template, it is possible to study functional connectivity between homotopic regions by simply flipping the x axis in stereotaxic space, e.g. [39]. The s-MRI scans were again corrected for intensity non-uniformities in stereotaxic space, this time restricted to the template brain mask. An individual brain mask was extracted a second time on this improved image [38] and combined with template priors. An iterative nonlinear registration was estimated between the linearly registered s-MRI and the template space, restricted to the brain mask [40]. A final brain mask of the T1 image in native space was extracted from the template brain mask by inverting the linear and non-linear transformation. This final mask was used for registration between rs-fMRI and sMRI data (see below). *Shared s-MRI outputs include*: non-uniformity corrected T1 volumes in native and stereotaxic space (after linear or non-linear transformations) at 1 mm isotropic resolution and brain masks in all spaces, in compressed NIFTI format (.nii.gz), as well as the linear and non-linear transformations from native to template space, as.xfm MINC files.

#### 3.3.2 Functional processing

*Preprocessing*. The NIAK rs-fMRI pipeline involved removing the first three volumes to allow for magnetization to reach equilibrium, site-specific slice timing correction to the middle slice, and estimating the parameters of a rigid-body motion between each time frame and the median volume of a run, followed by spatial resampling across frames. The fMRI time series were then corrected from slow time drifts (high-pass filter with a 0.01 Hz cut-off, using a discrete cosines transform) and physiological noise using an automated labeling of noise components in an individual independent component analysis, ICA [41]. Finally, the median volume of one selected fMRI run for each subject was coregistered (restricted to the brain) with the corresponding s-MRI scan using Minctracc [40]. The rs-fMRI to s-MRI transform and s-MRI to template (non-linear) transform were combined to resample the rs-fMRI volumes into MNI space at a 3 mm isotropic resolution and the results were spatially smoothed with a 6 mm FWHM Gaussian filter. *Shared rs-fMRI outputs include*: denoised rs-fMRI volumes in MNI space (compressed 4D NIfTIs, nii.gz), the mean/standard deviation rs-fMRI volumes and brain mask in native and template space (.nii.gz), six parameter head motion traces (HDF5.mat files) as well as individual ICA reports (.pdf).

*Time series for functionally defined regions*. A region-growing algorithm [42] based on the iterative merging of mutual-nearest-neighbours was implemented to generate functional brain parcellations. The spatial dimension was selected arbitrarily by specifying the size where the growing process should stop, measured in mm^3^. Two parameters (1000 mm^3^ and 330 mm^3^) were selected, resulting in the ROI1000 and ROI3000 parcellations, which include roughly 1000 and 3000 ROIs covering the grey matter, respectively. The region growing was applied on the time series concatenated across all participant’s rs-fMRI data (after correction to zero mean and unit variance) from the KKI site (training data only). The homogeneity of regions was thus maximized on average for all subjects, and the regions were identical for all subjects. To limit the amount of memory required by the region-growing procedure, it was applied seperatedly in each of the 116 areas of the AAL template [18]. The average time series for each ROI were extracted for both parcellations and are distributed in individual HDF5 (.mat) files. The R0I1000 and R0I3000 parcellations are also available as compressed 3D NIfTI files (.nii.gz).

#### 3.3.3 Quality control

Outputs of the NIAK pipeline were subjected to a careful visual inspection and the QC reports, along with head motion statistics, are available on the NIAK description page ^21^. Estimates of the maximum motion (translation and rotation) between consecutive functional volumes for each rs-fMRI dataset were inspected to categorize the datasets as containing minimal (< 1mm or degree), moderate (2 to 3 mm or degrees) or severe motion (>3 mm or degrees). The individual brain registration of the NIAK pipeline were visually inspected using online QC reports^22^, similar to those generated for the Athena pipeline. When substandard registration outcomes were identified, a parameter controlling the non-uniformity correction of the s-MRI was adjusted and the analysis was repeated until the coregistration results were satisfactory. Satisfactory results could not be achived with some datasets and have been indicated as “Fail” for QC (5.2% failure rate) in the.tsv spreadsheet including QC assessments, available on NITRC^23^.

## 4. Usage recommendations

In line with the original purpose of the ADHD-200 Global competition, most of the publications that have used the ADHD-200 Preprocessed initiative data have proposed new machine learning techniques for predicting ADHD diagnosis or subtype. The most frequently used derivatives have been Athena preprocessed fMRI volumes [e.g. 43, 44] and Athena regional time series [e.g. 45, 46]. Works have used Burner pipeline data both in isolation [e.g. 47–49] or in combination with the Athena functional derivatives [e.g. 50–52]. A fewer number of publications have also used Athena ICN, ReHo and fALFF maps [e.g. 53] and the NIAK high resolution (either R0I1000 or R0I3000) regional time series in their main analysis [e.g. 54, 55]. Researchers have also found value in this initiative beyond the processed data, such as the CC200 or CC400 functional brain parcellations [e.g. 56, 57] and the Athena processing scripts [.e.g 58]. The three PhD dissertations [59–61] and three master’s theses [62–64] that have used this resource have all focused on disease state prediction and data dimensionality reduction techniques.

To the best of our knowledge, there are no published comparisons between the results generated by the Athena and NIAK pipeline. The two pipelines conceptually implement very similar steps, with key operations including the non-linear volumetric registration of individual structural MRIs in two different variants of the MNI space, and the registration of individual EPI and T1 images. The most striking differences between the pipelines are in the software that they used and the opinions that drove the parameter choices. The Athena implemented nuisance variable regression [14, 15] (motion parameters, WM and GM signals), while NIAK implemented an automated labeling of structured noise in a spatial ICA [41]. The Athena strategy is fairly standard in the rs-fMRI community, while ICA-based noise attenuation techniques are less common. Athena provided time series for a variety of low-dimension anatomical and functional brain parcellations, whereas NIAK favored much higher-resolution (several thousands) brain parcels. There are also differences in the type of data that each pipeline offers, Athena provides a wide variety of statistical derivatives calculated from both the functional and structural data, whereas Burner provided structural derivatives only, and NIAK primarily focused on function. Beyond personal ideology, the availability of particular features, or allegiances to certain software tools there is no clear reason to prefer one pipeline over the other. It has recently become clear that even subtle variations of processing environments can impact the end-results of processing pipelines [65], even when the same processing is replicated. The variety of data available through this initiative enables researchers to compare the robustness of their tools or analysis results to different processing choices and derivatives.

The ADHD-200 Preprocessed sample also provides ample opportunity for analyses besides the identification of rs-fMRI based biomarkers of ADHD [66]. With 585 TDC participants between the ages of 7 and 26, and the inclusion of intelligence measures, the ADHD-200 Preprocessed is a valuable resource for mapping developmental trajectories [67–69] and other sources of inter-individual variation [70]. Perhaps most exciting are new methods that cluster individuals based on connectivity profiles [71, 72], which are providing new hope for using neuroimaging data to parse the heterogeneity within mental health disorders [73]. One of the outstanding needs for neuroimaging, and connectomics in particular, is the development and validation of new analytical tools and processing strategies [66, 74, 75]. In the service of this aim, the ADHD-200 Preprocessed repository has the necessary components to become a benchmark dataset for evaluating new tools as they are proposed.

The two biggest challenges for using the ADHD-200 Preprocessed data are head motion [76–81] and inter-site variation in the acquisition equipment, parameters, and experimental procedures [82, 83]. A variety of different approaches have been proposed for addressing head motion in hyperkinetic populations [76, 84], and in the ADHD-200 Sample in particular [79], that should be considered when analyzing the data. At the very least, some statistic that characterizes individual motion (such as root mean square deviation [85]) should be included as a nuisance regressor in the group-level model [78, 80]. Differences in the manner in which data was collected at each site can introduce additive and multiplicative effects (batch effects) to the data, which may obscure the underlying biological signal [82, 83]. Including a regressor for acquisition protocol (see Table 2 and Table 3 for a summary of the different protocols), the average pairwise correlation between all regions in the brain (GCOR) [86], or the whole-brain average of the feature under inquiry [83], have all been shown to be effective for dealing with inter-site variation.

## 5. Discussion and conclusions

The ADHD-200 Preprocessed initiative was successful in terms of its primary objectives: the derivatives shared in the repository were effectively used by many researchers during and after the ADHD-200 Global Competition, with over 10,500 downloads by more than 600 users, as well as 52 resulting publications [34, 43–58, 82, 87–120], 3 PhD theses [59–61], 3 master’s dissertations [62–64], and 1 patent [121] derived from the release in just over three years. Further publications are either in press or under review, and an updated list of publications will be maintained in a public Mendeley group^24^. Although there was clearly a peak in usage around the ADHD-200 Global competition, there has been a sustained amount of downloads and publications since then, see Figure 5, which we take as a demonstration of a long-term interest from the community in this resource.

**Figure 5:**
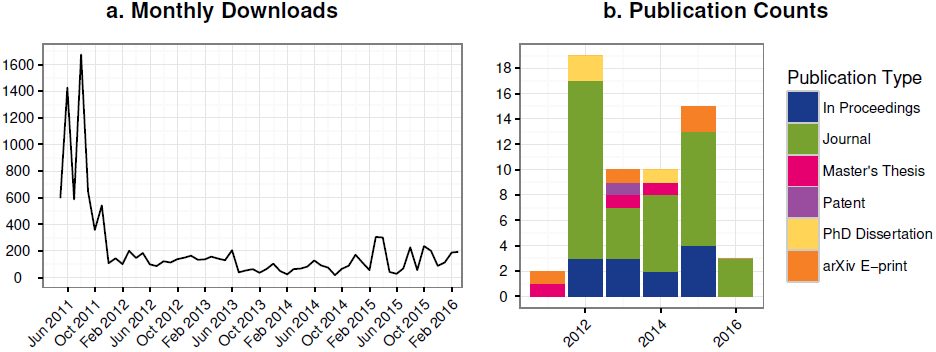
Statistics on download and citations of the ADHD200 preprocessed initiative.

The ADHD-200 preprocessed has helped to expand the boundaries of the traditional neuroimaging community, with several publications in core engineering and statistics journals that do not routinely feature neuroimaging applications, such as Statistica Sinica [55] or the Journal of the American Statistical Association [104], amongst others [e.g. 43, 47–50, 54, 89–91, 99–102, 113, 114]. In particular, the winning team of the ADHD-200 Global Competition was based at the Johns Hopkins Biostatistics Department and used ADHD-200 Preprocessed to develop their diagnostic algorithm [51]. We also found interesting that a handful of methodological publications used the ADHD-200 Preprocessed sample as one application in a series of benchmarks that could be as varied as positron emission tomography in Alzheimer’s disease [55], the Yeast gene regulatory network [102] or gene expression in brain tissues of patients with HIV-1 associated neurocognitive disorders [91]. This suggests that fully processed, easy-to-access imaging samples could help validate general-purpose methods on a wider scope of applications.

The impact of the ADHD-200 Preprocessed repository demonstrated the need for reducing computational barriers to participation in discovery neuroscience, including but not limited to machine learning competitions based on neuroimaging data. While the ADHD-200 Preprocessed initiative will have a long-term impact on that need, we believe that a much larger-scale effort will be necessary to unlock the full potential of openly shared neuroimaging data in the service of accelerating neuroscience research. In line with the grassroots, open science ethos of the Neurobureau, new contributors interested in sharing derivatives of ADHD-200 or other open imaging data repository can contact us via our web-based forum^25^. We are actively seeking new contributions to the Preprocessed Connectomes Project, notably during Brainhack events[122]. An important current project is the ABIDE preprocessed initiative^26^ [69]. This new resource, still under development, will include 16 different processing strategies for rs-fMRI processing, implemented across 4 different software packages, and 2 different pipelines for structural MRI processing. The release will feature an harmonized organization of processed derivatives across packages and extensive quality control. We are planning to continue to expand this line on work on other data sources in the near future. Our hope is that ADHD-200 Preprocessed and future related efforts will critically help fMRI researchers to identify optimal analytical paths for a given task.

## Acknowldegements

The authors would like to thank the ADHD-200 Consortium for assembling and sharing the ADHD-200 Sample and for hosting the ADHD-200 Global competition. RCC would like to thank the developers of AFNI and FSL, and the Advanced Research Computing at Virginia Tech for computational support. The scripts used to calculate ReHo were generously provided by Dr. Xinian Zuo. CC would like to thank the developers of SPM. PB would like to thank the developers of MINC tools, the CIVET pipeline, the CBRAIN computational infrastructure, and the computational resources provided by Compute Canada^27^ and CLUMEQ^28^, which is funded in part by NSERC (MRS), FQRNT, and McGill University.

## Footnotes

1 See http://www.neurobureau.org/mission-statement/ for the full mission statement. The Neurobureau is a non-profit organization registered in Germany

2 http://www.nitrc.org/frs/?group_id=383

3 http://fcon_1000.projects.nitrc.org/indi/ADHD-200/

4 http://www.nitrc.org/forum/forum.php?forum_id=2046

5 http://www.nitrc.org/forum/forum.php?forum_id=1735.

6 Further information regarding excluded data can be found at the respective pipeline wiki page: Athena: http://www.nitrc.org/plugins/mwiki/index.php/neurobureau:AthenaPipeline#Excluded_Data; Burner: http://www.nitrc.org/plugins/mwiki/index.php/neurobureau:BurnerPipeline; NIAK: http://www.nitrc.org/plugins/mwiki/index.php/neurobureau:NIAKPipeline.

7 http://www.nitrc.org/plugins/mwiki/index.php/neurobureau:AthenaPipeline

8 http://www.arc.vt.edu/

9 https://github.com/preprocessed-connectomes-project/adhd200_athena_scripts

10 http://www.fz-juelich.de/inm/inm-1/EN/Forschung/_docs/SPMAnatomyToolbox/SPMAnatomyToolbox_node.html

11 http://www.cyceron.fr/index.php/en/plateforme-en/freeware

12 http://preprocessed-connectomes-project.org/adhd200_visual_qc_athena/

13 https://www.nitrc.org/frs/download.php/9024/adhd200_preprocessed_phenotypics.tsv

14 http://www.nitrc.org/plugins/mwiki/index.php/neurobureau:BurnerPipeline

15 http://www.nitrc.org/plugins/mwiki/index.php/neurobureau:NIAKPipeline

16 http://en.wikibooks.org/wiki/MINC

17 http://www.nitrc.org/frs/?group_id=411

18 https://github.com/SIMEXP/niak

19 https://github.com/preprocessed-connectomes-project/adhd200_niak_scripts

20 http://psom.simexp-lab.org/how_to_use_psom.html

21 http://www.nitrc.org/plugins/mwiki/index.php/neurobureau:NIAKPipeline#Quality_control_of_the_preprocessing_-_Training_dataset

22 http://preprocessed-connectomes-project.org/adhd200_visual_qc_niak/

23 https://www.nitrc.org/frs/download.php/9024/adhd200_preprocessed_phenotypics.tsv

24 https://www.mendeley.com/groups/4198361/adhd-200-preprocessed/

25 http://www.nitrc.org/forum/?group_id=383

26 http://preprocessed-connectomes-project.org/abide/

27 https://computecanada.org/

28 http://www.clumeq.mcgill.ca/

